## Supplement for "Neural representation of abstract task structure during generalization"

| | Initial Training (phase 1) | | | | New Category Training (phase 2) | Generalization (phase 3) | |
|---|---|---|---|---|---|---|---|
| | Initial Training 1 | Initial Training 2 | Initial Training 3 | Initial training 4 (reminder) | New Category Training | Generalization (mini-blocked) | Generalization (mixed) |
| **Contexts** | A1, B1, C1 | A2, B2, C2 | A3, B3, C3 | A1, B1, C1<br>A2, B2, C2<br>A3, B3, C3 | A3, B3, C3 | A1, B1, C1<br>A2, B2, C2 | A1, B1, C1<br>A2, B2, C2 |
| **Item categories** | Hands, foods, leaves | Hands, foods, leaves | Hands, foods, leaves | Hands, foods, leaves | Faces, animals, objects | Faces, animals, objects | Faces, animals, objects |
| **Session 1** | 2 blocks x 54 trials | 2 blocks x 54 trials | 2 blocks x 54 trials | 1 block x 216 trials | 1 block x 90 trials | 1 block x 180 trials | - |
| **Session 2** | 2 blocks x 54 trials | 2 blocks x 54 trials | 2 blocks x 54 trials | 1 block x 216 trials | 1 block x 90 trials | 1 block x 180 trials | 3 blocks x 180 trials |
| **Session 3** | 1 block x 54 trials | 1 block x 54 trials | 1 block x 54 trials | 1 block x 216 trials | 1 block x 90 trials | 1 block x 180 trials | 3 blocks x 180 trials |

**Figure 1 – figure supplement 1.** Schematic showing design of whole experiment over multiple sessions. Note that contexts seen in new category training and held out of generalization were not necessarily from Initial Training 3, but these are shown only as an example here. The mixed generalization phase did not take place in session 1. Only participants who passed the accuracy criterion in the generalization phase in session 1 completed the mixed generalization phase in the scanner, while participants who failed to meet this criterion completed this phase behaviorally. All other sections of the experiment took place outside of the scanner.

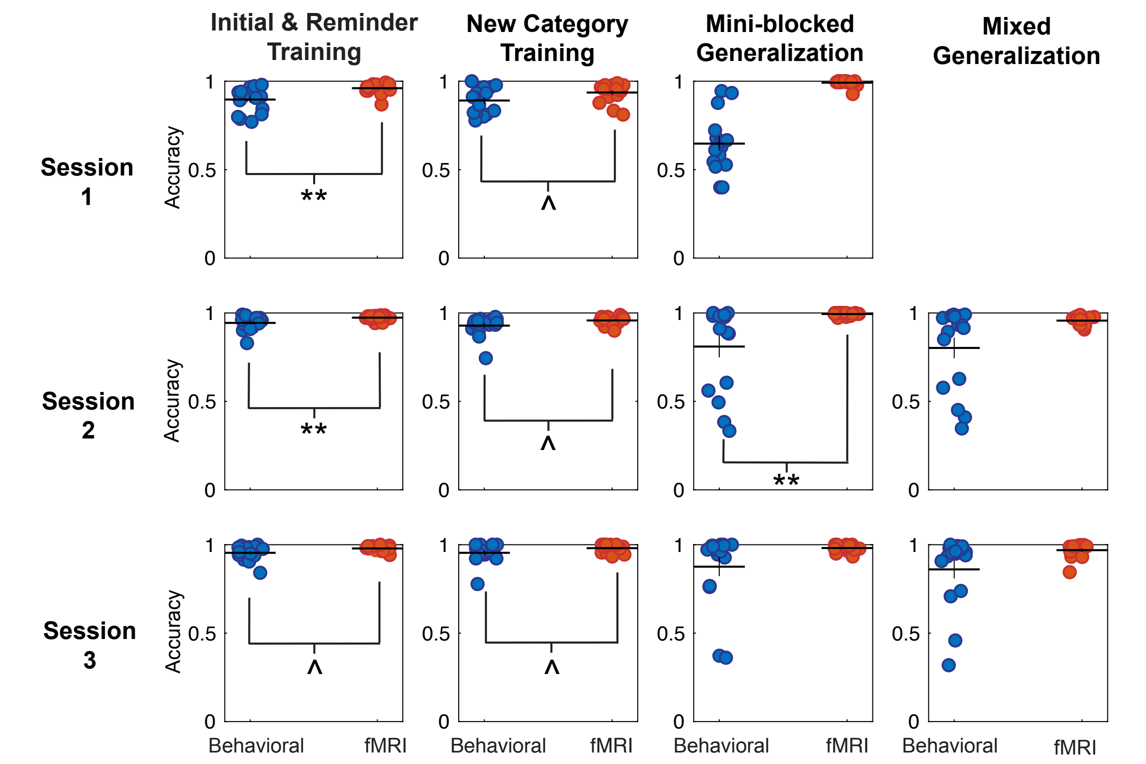

**Figure 2 - figure supplement 1.** Accuracy in all sessions of the experiment for behavioral and fMRI groups. No statistical comparison between groups was made in the mini-blocked generalization phase of session 1 as accuracy in this phase was used to define these groups. Each point represents a single participant, horizontal bars represent the group mean and vertical bars indicate the standard error of the mean. ** $P < 0.01$, * $P < 0.05$, ^ $P < 0.1$, Wilcoxon rank sum tests between groups.

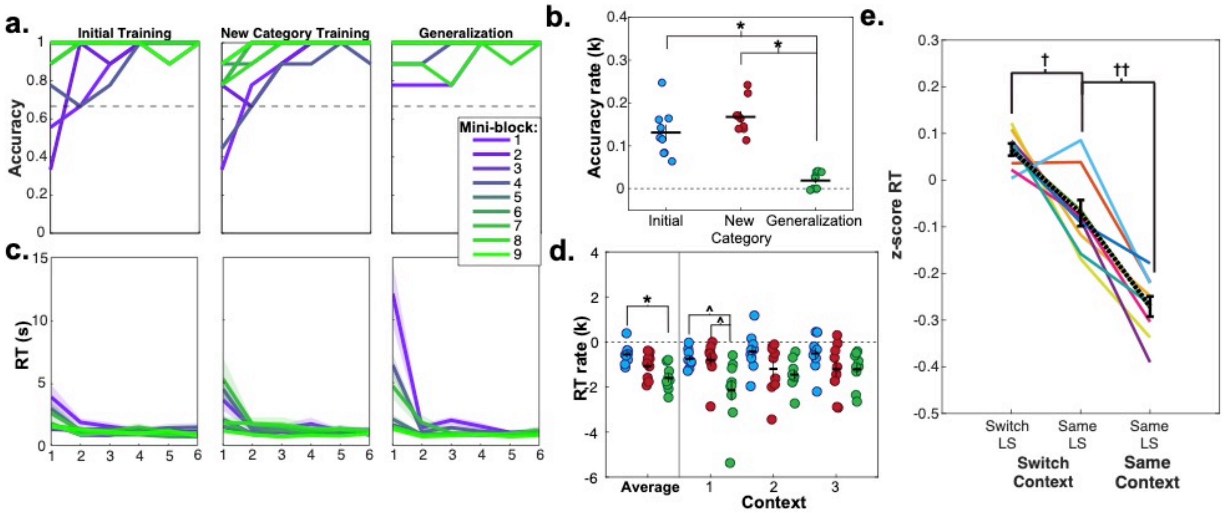

**Figure 2 – figure supplement 2.** Learning curves and repetition effects from training and generalization phases for behavioral participants who passed the accuracy criterion for generalization in session 2 (N = 9). **a**. Mean accuracy across participants for first six trials of the first nine mini-blocks of initial and new category training, as well as blocked generalization phase. Dashed line indicates chance-level performance. **b.** Estimated exponential rate of change in mean accuracy in the first six trials for all mini-blocks in each phase, across participants. **c.** Mean reaction times (RT) for each trial in each of the first nine mini-blocks for each of these phases. Shaded area represents standard error of the mean (SEM). Mean and standard error were calculated from the log-transformed RTs. **d.** Estimated exponential rate of change in RT for first six trials within each phase for the first mini-blocks within the first three contexts in presentation order. For panels **b** and **d**, each dot represents a single participant, horizontal and vertical bars represent the mean and SEM, respectively. * $P < 0.05$, Wilcoxon signed rank test, corrected for multiple comparisons, ^ $P < 0.05$, uncorrected. **e.** RTs for trials from the pseudo-randomized generalization phase from sessions 2 and 3 during fMRI scanning where latent states remained the same or switched from the previous trial, while the context switched or stayed the same. RTs have been log-transformed and z-scored within session for each participant. Each line represents a participant, black dashed line indicates mean and error bars indicate SEM. †$P < 0.05$, ††$P < 0.01$, repeated-measures t-test.

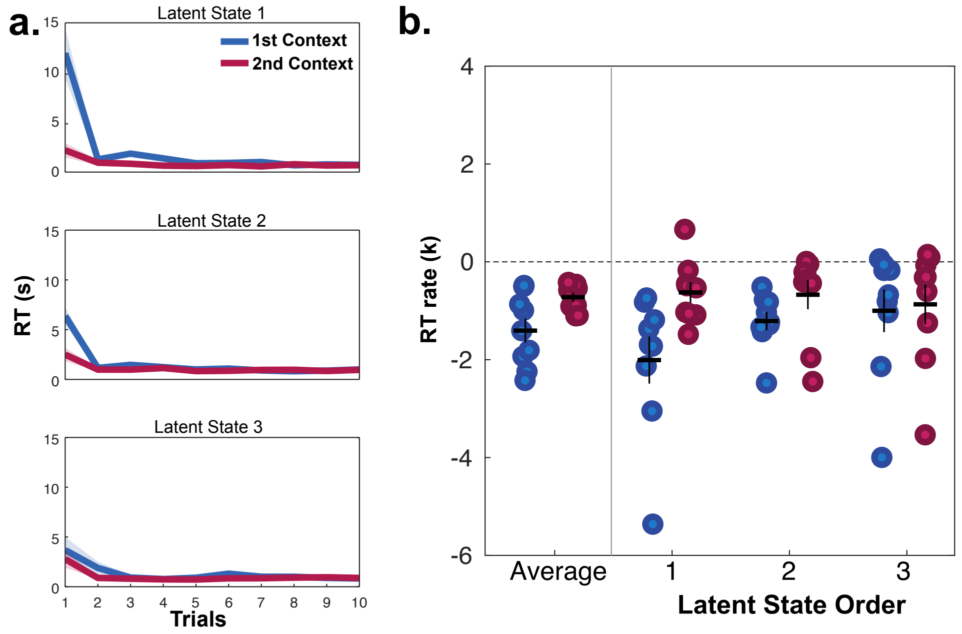

**Figure 3 – figure supplement 1.** Reaction times (RT) for participants in the behavioral group who passed the generalization criterion in session 2 (N = 9) during mini-blocks during generalization in session 2, organized by the presentation of latent states (LS 1, 2, 3) and contexts ($1^{st}$ and $2^{nd}$). Participants were significantly slower on the first trial of mini-blocks when encountering the first context in a latent state compared to the second context in the same latent state), but not in subsequent trials **a**. RT curves for mini-blocks organized by the order of latent states and contexts encountered. Lines indicate means, shaded area indicates the SEM. **b.** Rates of exponential functions fit to these RT curves. Each dot represents a participant. Horizontal bars indicate the mean and vertical bars the SEM.

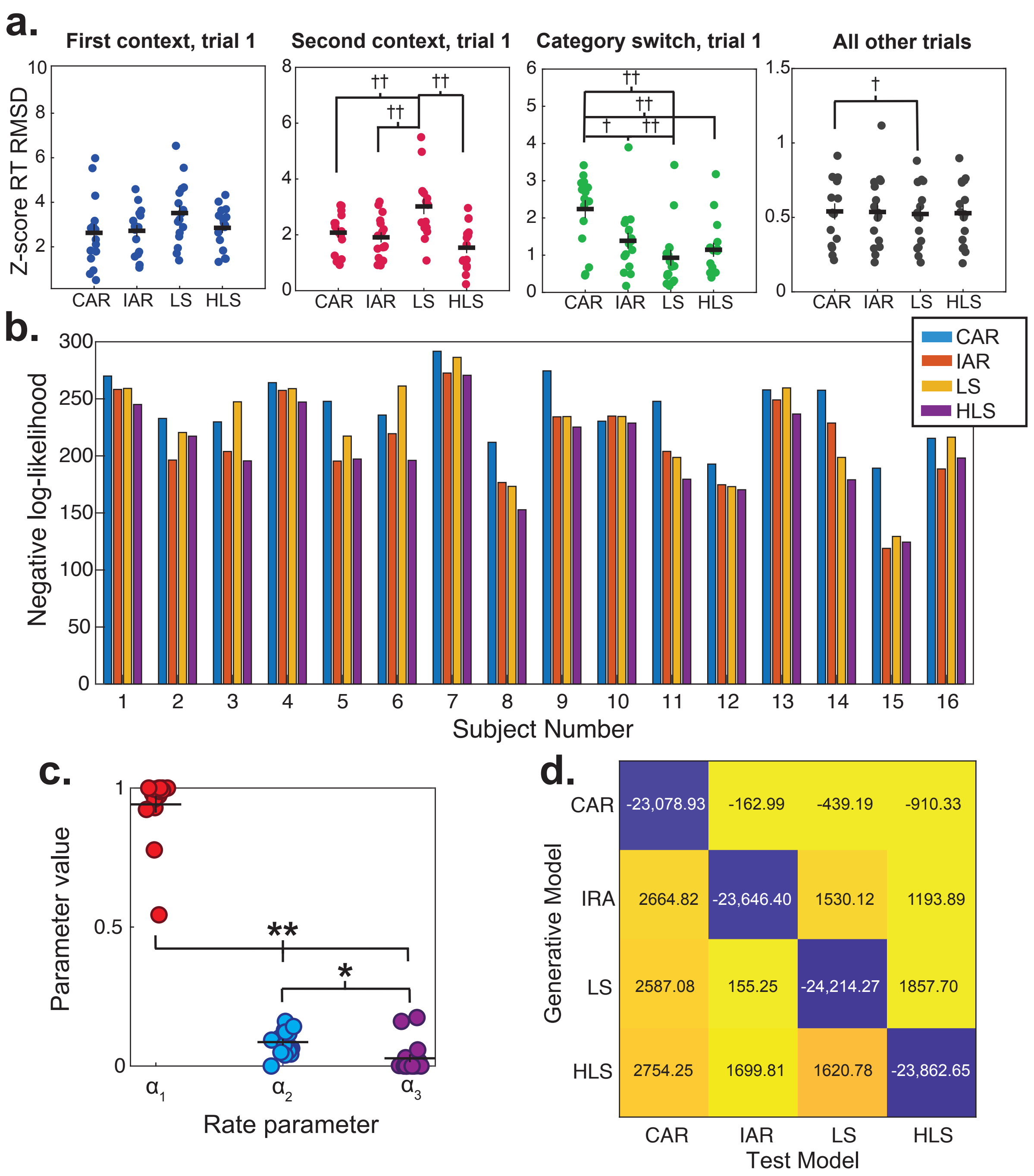

**Figure 4 – figure supplement 1.** Supplementary analyses for computational modeling of reaction times (RTs). Data are from session 2 of the mini-blocked generalization phase for participants in the fMRI group. **a.** Root-mean squared deviation between data simulated from each model and participants' z-scored reaction times for different trial-types. † $P < 0.05$, †† $P < 0.01$, repeated-measures t-tests, Bonferroni corrected for multiple comparisons. **b.** Negative log-likelihoods for each participant in the fMRI group for conjunctive associative retrieval (CAR), independent associative retrieval (IAR), latent state (LS) and hierarchical latent state (HLS) models. Lower values indicate a better fit. **c.** Values for three rate parameters governing rate of activation change in nodes due to direct ($\alpha_1$), mediated ($\alpha_2$) and incidental retrieval ($\alpha_3$). **d.** Cross model comparison using best fitting parameters from 16 participants to generate simulated data, tested with alternative. Values indicate negative log-likelihood of model fits. ** $P = 0.001$, * $P < 0.05$, Wilcoxon signed rank test, Bonferroni corrected for multiple comparisons.

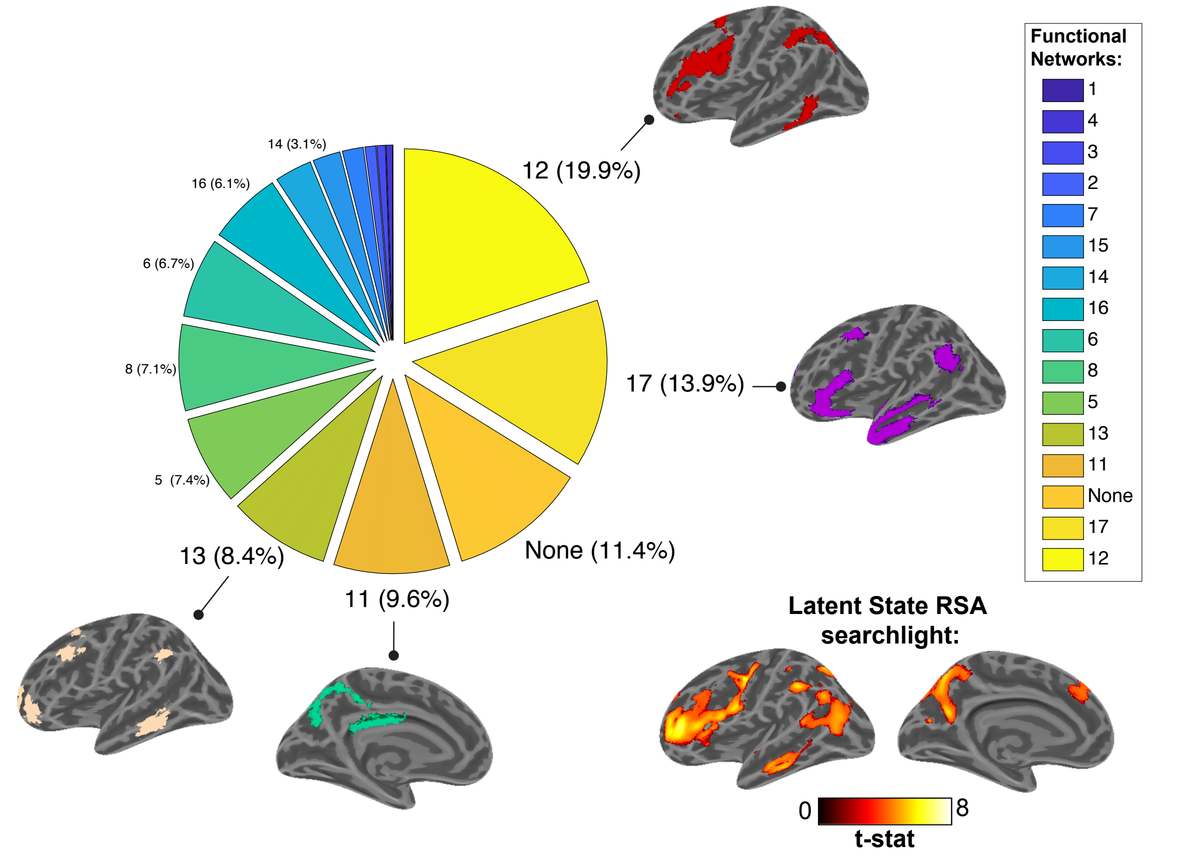

**Figure 5 – figure supplement 1.** Percentage overlap of latent state statistical map from Figure 5a with 17 network parcellation of functional connectivity networks from Yeo et al. (2011), based on the data of 1000 participants. Numbers correspond to network identifications from the same study with percentage of voxels within each network shown in parentheses and represented by the pie chart. Surface renderings of the three networks that included the largest proportion of voxels are shown on an inflated cortical surface. Bottom right-hand corner shows the t-statistic map for the latent state regressor from the representational similarity analysis searchlight for comparison, also shown in Figure 5a.

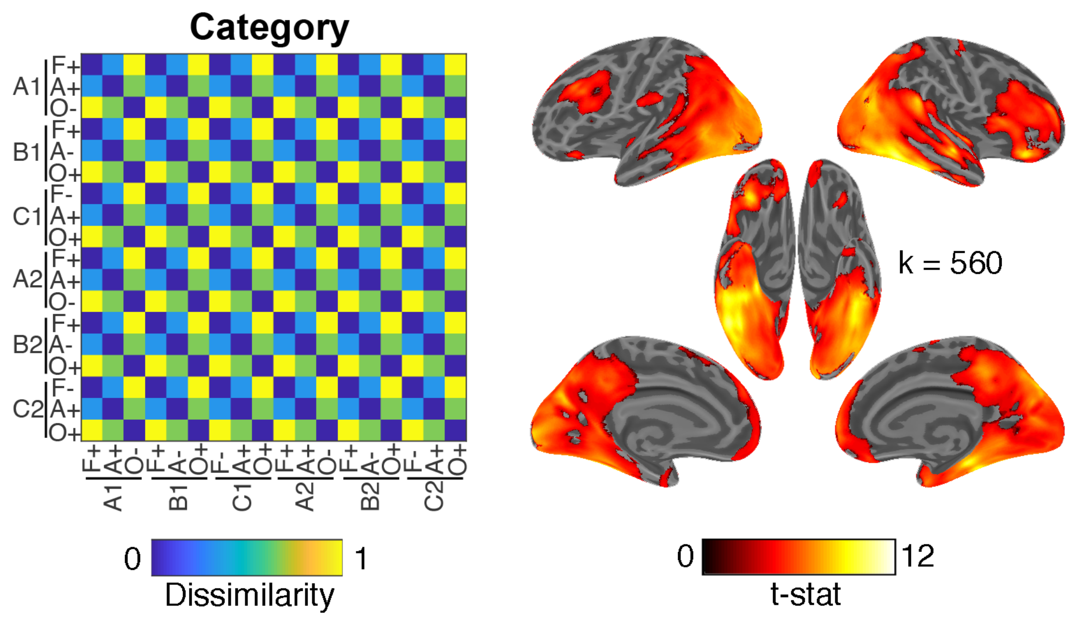

**Figure 5 – figure supplement 2.** Whole brain searchlight analysis for image categories. Left panel shows hypothesis representational dissimilarity matrix for categories (based on animacy). Right panel shows the cluster corrected t-statistic map from a searchlight analysis projected onto an inflated cortical surface. This statistical map statistical map was thresholded with a cluster forming threshold of $P < 0.001$ and corrected for multiple comparisons using cluster-based permutation tests at $P < 0.05$. The cluster extent threshold from this test is given by value of k.

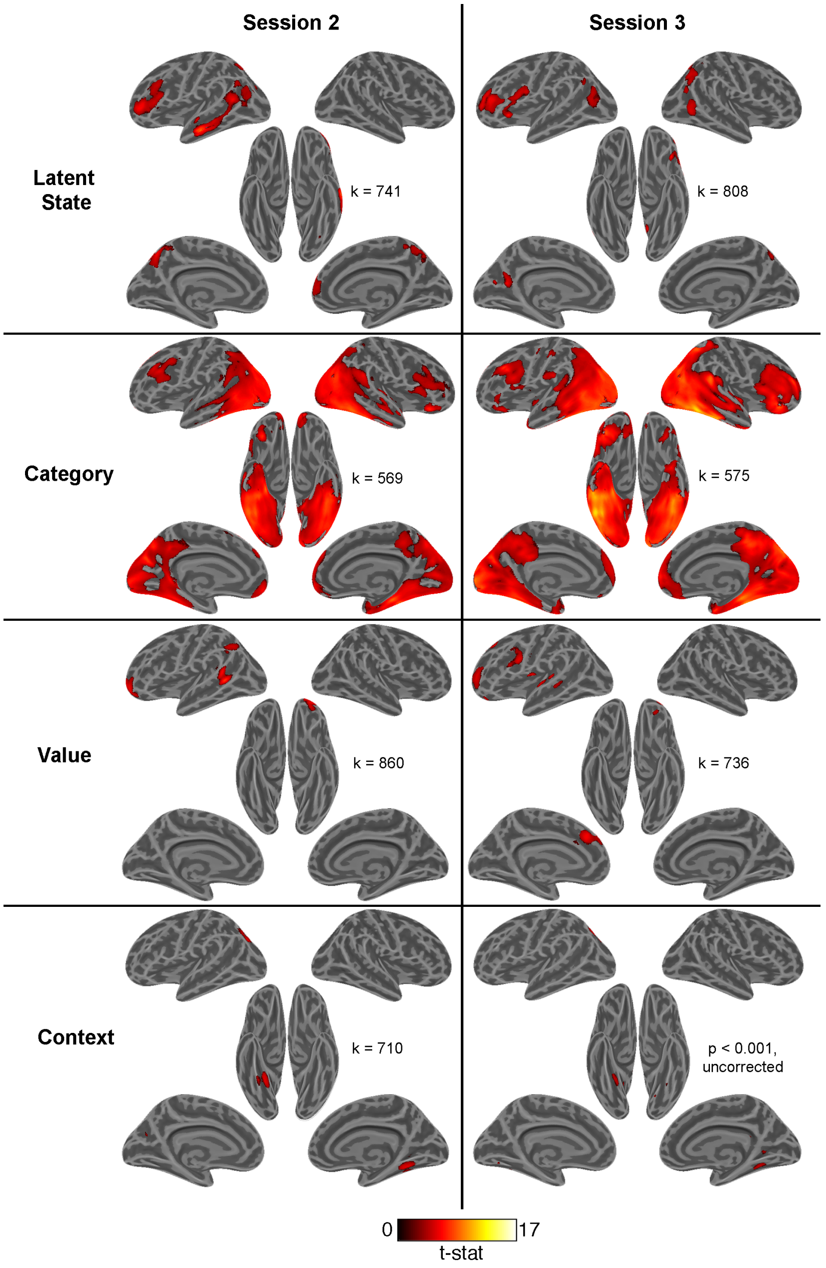

**Figure 5 – figure supplement 3.** Statistical maps for main effects of multiple regression model from searchlight representational similarity analyses (RSAs) estimated separately for data from sessions 2 and 3 and projected onto inflated cortical surfaces. All statistical maps were thresholded with a cluster forming threshold of $P < 0.001$ and corrected for multiple comparisons using permutation tests to find a cluster extent threshold (k) at $P < 0.05$, except for the context effects in session 3 where no clusters passed this threshold (these data are shown at $P < 0.001$, uncorrected).

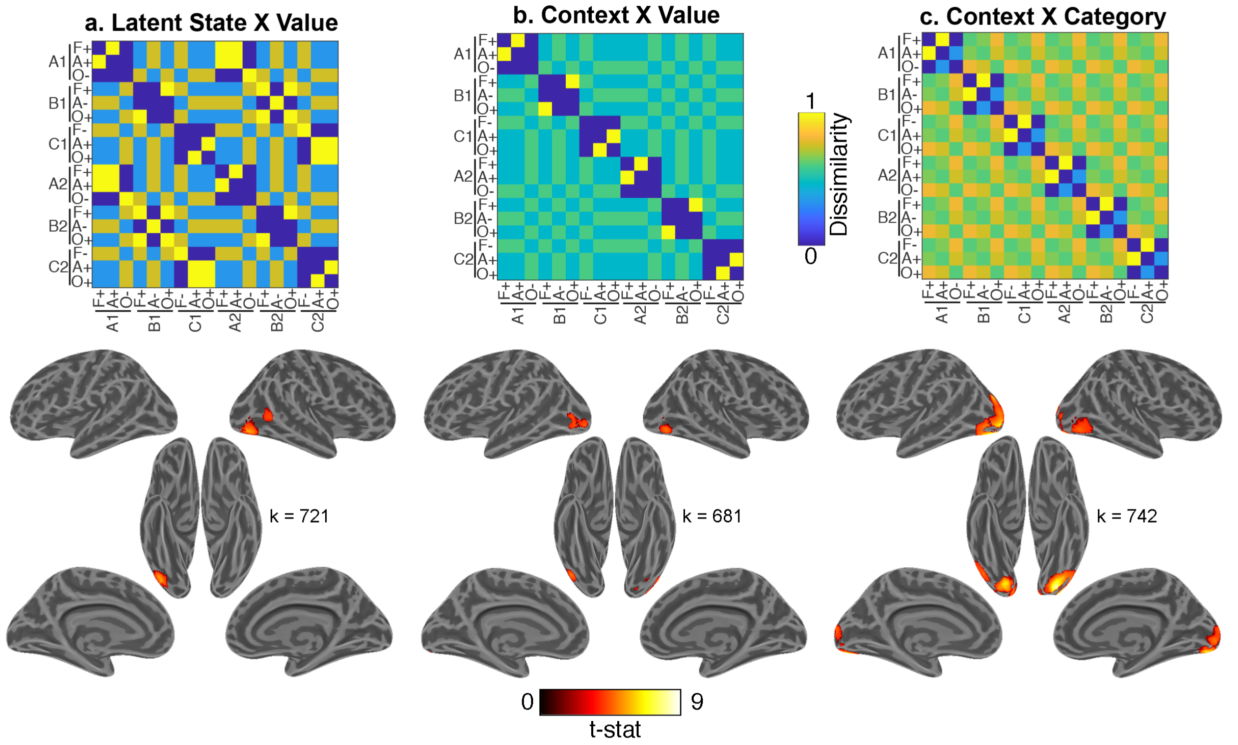

**Figure 5 – figure supplement 4.** Whole-brain searchlight analyses for interaction terms. **a.** Results showing where pattern activity was modulated by the negative interaction of latent state and value. Nuisance regressors for interactions between contexts and **b.** value and **c.** categories. All maps are defined with a cluster forming threshold of $P < 0.001$ and corrected for multiple comparisons with permutation tests for defining a cluster extent threshold at $P < 0.05$.

35

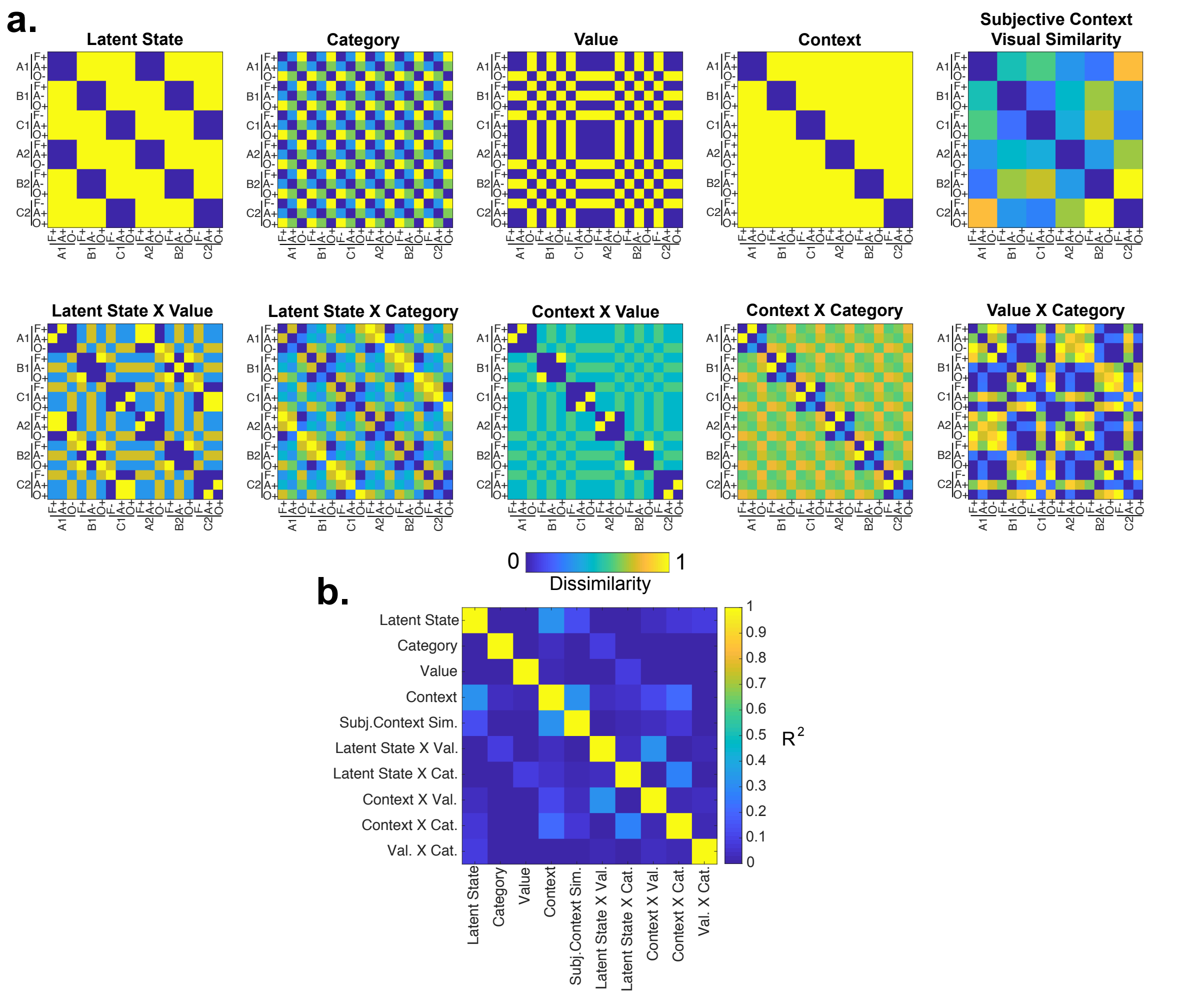

**Figure 5 – figure supplement 5.** Hypothesis representational dissimilarity matrices (RDMs). **a.** All hypothesis RDMs included in the multiple regression analysis for searchlight and ROI-based representational similarity analyses (RSA). Color bar indicates dissimilarity. A, B, C refer to distinct latent states. A1, B1, C1 and A2, B2, C2 refer to distinct contexts that belong to each of those latent states. F, Faces; A, Animals; O, Objects. +, positive value; -, negative value. **b.** Variance shared ($R^2$) between hypothesis RDMs.

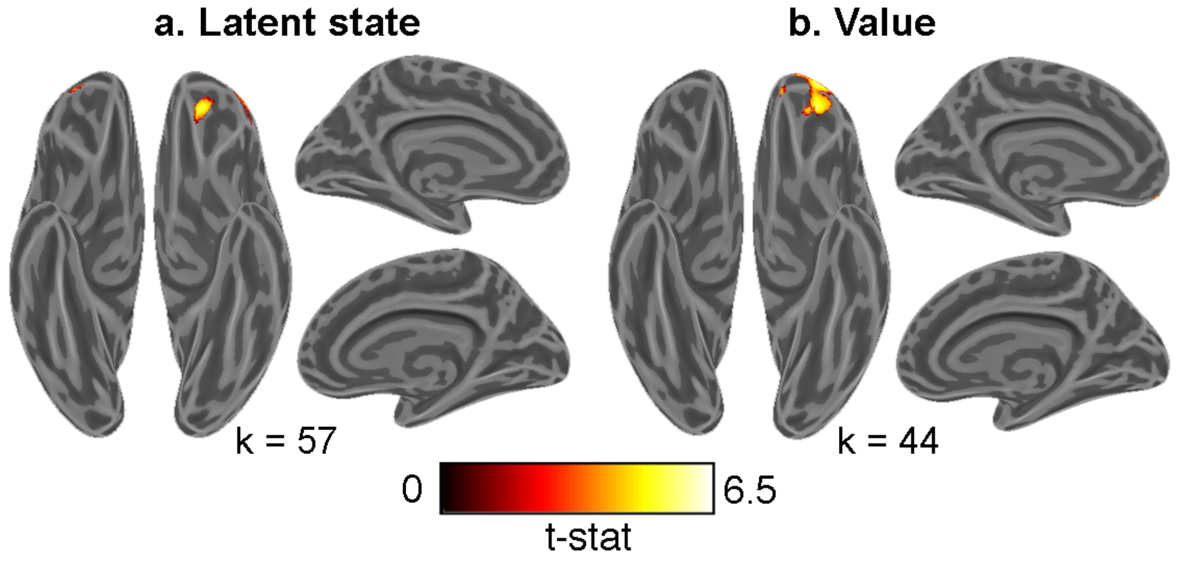

**Figure 6 – figure supplement 1.** Results of representational similarity analysis searchlight results with explicit mask in orbitofrontal cortex. Maps of t-statistics have been projected onto an inflated cortical surface to show effects for **a.** Latent states and **b.** Value. Both maps were defined with a cluster forming threshold of $P < 0.001$ and corrected for multiple comparisons with permutation tests for defining a cluster extent threshold at $P < 0.05$. The cluster extent threshold is given by the value of k.

**Supplementary Table 1. Activations passing permutation-based cluster correction for whole-brain representational similarity analysis**

| Region (AAL2) | MNI Coordinates | | | Number of voxels | Peak *t*-value |
|---|---|---|---|---|---|
| | x | y | z | | |
| **Latent state** (k = 760) | | | | | |
| Left inferior frontal gyrus, triangularis | -34.5 | 42 | 4.5 | 11206 | 7.49 |
| Left precentral gyrus | -51 | -4.5 | 49.5 | | 7.25 |
| Left precentral gyrus | -55.5 | 6 | 22.5 | | 5.80 |
| Right postcentral gyrus | 43.5 | -30 | 42 | 1265 | 7.16 |
| Right inferior parietal gyrus | 58.5 | -45 | 49.5 | | 4.22 |
| Left middle temporal gyrus | -63 | -13.5 | -10.5 | 12763 | 6.79 |
| Right precuneus | 6 | -72 | 51 | | 6.53 |
| Left calcarine fissure | -9 | -63 | 18 | | 6.14 |
| Right insula | 34.5 | 22.5 | 12 | 1807 | 6.11 |
| Right middle frontal gyrus | 40.5 | 55.5 | 9 | | 5.57 |
| Right middle frontal gyrus | 45 | 30 | 34.5 | 1807 | 3.97 |
| Left inferior parietal gyrus | -49.5 | -37.5 | 45 | 1334 | 6.07 |
| **Context** (k = 726) | | | | | |
| Right fusiform gyrus | 31.5 | -48 | -6 | 1386 | 6.04 |
| Left fusiform gyrus | -25.5 | -64.5 | -16.5 | 1102 | 5.30 |
| Left middle occipital gyrus | -22.5 | -63 | 39 | 948 | 5.11 |
| **Value** (k = 760) | | | | | |
| Left superior frontal gyrus, dorsolateral | -27 | 58.5 | -1.5 | 2473 | 6.01 |
| Left anterior orbital gyrus | -27 | 42 | -13.5 | | 5.75 |
| Left middle temporal gyrus | -52.5 | -49.5 | 10.5 | 2140 | 4.85 |
| Left angular gyrus | -45 | -52.5 | 34.5 | | 4.44 |
| Left middle occipital gyrus | -27 | -63 | 37.5 | | 4.41 |
| Left superior frontal gyrus, dorsolateral | -15 | 46.5 | 40.5 | 808 | 4.66 |
| **Category** (k = 560) | | | | | |
| Vermis 7 | 0 | -76.5 | -19.5 | 183997 | 11.28 |
| Right inferior temporal gyrus | 46.5 | -57 | -13.5 | | 10.69 |
| Right Cerebelum 4 5 | 25.5 | -33 | -24 | | 10.48 |
| Left posterior orbital gyrus | -36 | 25.5 | -22.5 | 1962 | 6.07 |
| Left fusiform gyrus | -31.5 | -3 | -40.5 | | 4.54 |
| Left amygdala | -27 | -1.5 | -18 | | 4.45 |
| Right supplementary motor area | 15 | -10.5 | 67.5 | 563 | 5.75 |

| | | | | | |
|---|---|---|---|---|---|
| **Latent State X Value x -1**<br>(k = 721) | | | | | |
| Right inferior occipital gyrus | 48 | -76.5 | -13.5 | 2139 | 6.54 |
| **Context X Value**<br>(k = 681) | | | | | |
| Right inferior occipital gyrus | 48 | -78 | -7.5 | 865 | 5.30 |
| Left middle occipital gyrus | -48 | -79.5 | 1.5 | 1015 | 4.83 |
| Left occipital gyrus | 48 | -78 | -7.5 | | 4.59 |
| **Context X Category**<br>(k = 742) | | | | | |
| Left lingual gyrus | -27 | -88.5 | -18 | 8442 | 8.57 |
| Left middle occipital gyrus | -25.5 | -82.5 | 15 | | 6.63 |
| Right lingual gyrus | 21 | -82.5 | -10.5 | | 6.57 |
| Right inferior occipital gyrus | 48 | -78 | -7.5 | 1293 | 6.27 |
| Right middle temporal gyrus | 52.5 | -72 | 15 | | 4.44 |

All reported clusters were significant at the $P < 0.05$, corrected for multiple comparisons after peak thresholding at $P < 0.001$ and permutation-based cluster correction. The critical cluster extent threshold for each contrast is given by the value of k.

**Supplementary Table 2. Activations passing permutation-based cluster correction for representational similarity analysis constrained to orbitofrontal cortex region of interest**

| Region (AAL2) | MNI Coordinates | | | Number of voxels | Peak *t*-value |
|---|---|---|---|---|---|
| | x | y | z | | |
| **Value** (k = 57) | | | | | |
| Left superior frontal gyrus, dorsolateral | -27 | 58.5 | -1.5 | 996 | 6.01 |
| Left anterior orbital gyrus | -27 | 42 | -13.5 | 996 | 5.75 |
| Left gyrus rectus | -4.5 | 57 | -21 | 57 | 4.32 |
| **Latent state** (k = 44) | | | | | |
| Left medial orbital gyrus | -19.5 | 42 | -19.5 | 232 | 6.05 |
| Right middle frontal gyrus | 36 | 58.5 | -3 | 169 | 5.05 |
| Left middle frontal gyrus | -42 | 43.5 | -1.5 | 211 | 4.87 |

All reported clusters were significant at the $P < 0.05$, corrected for multiple comparisons after peak thresholding at $P < 0.001$ and permutation-based cluster correction within an explicit mask defining orbitofrontal cortex. The critical cluster extent threshold for each contrast is given by the value of k.

**Supplementary Table 3. Activations passing permutation-based cluster correction for univariate contrast of correct and erroneous responses**

| Region (AAL2) | MNI Coordinates | | | Number of voxels | Peak *t*-value |
|---|---|---|---|---|---|
| | x | y | z | | |
| **Correct > error** (k = 317) | | | | | |
| Right hippocampus | 40.5 | -22 | -17.5 | 576 | 4.56 |
| Left superior temporal gyrus | -61.5 | -26.5 | 5 | 22560 | 6.79 |
| Left medial orbital gyrus | -7.5 | 48.5 | -14.5 | 558 | 4.92 |
| Right middle temporal gyrus | 40.5 | -40 | -8.5 | 349 | 4.70 |

All reported clusters were significant at the $P < 0.05$, corrected for multiple comparisons after peak thresholding at $P < 0.001$ and permutation-based cluster correction. The critical cluster extent threshold for each contrast is given by the value of k.
